# Identification of a novel Nucleobase-Ascorbate Transporter family member in fish and amphibians

**DOI:** 10.1101/287870

**Authors:** Diogo Oliveira, André M. Machado, Tiago Cardoso, Mónica Lopes-Marques, L. Filipe C. Castro, Raquel Ruivo

## Abstract

Nucleobase-Ascorbate Transporter (NAT) family includes ascorbic acid, nucleobases and uric acid transporters: with a broad evolutionary distribution. In vertebrates, four members have been previously recognized, the ascorbate transporters Slc23a1 and Slc3a2, the nucleobase transporter Slc23a4 and an orphan transporter Slc23a3. Using phylogenetic and synteny analysis, we identify a fifth member of the vertebrate *slc23* complement (*slc23a5*), present in neopterygians (gars and teleosts) and amphibians, and clarify the evolutionary relationships between the novel gene and known *slc23* genes. Further comparative analysis puts forward uric acid as the preferred substrate for Slc23a5. Gene expression quantification, using available transcriptomic data, suggests kidney and testis as major expression sites in *Xenopus tropicalis* (western clawed frog) and *Danio rerio* (zebrafish). Additional expression in brain was detected in *D. rerio*, while in the Neoteleostei *Oryzias latipes* (medaka) *slc23a5* expression is restricted to brain. The biological relevance of the retention of an extra transporter in fish and amphibians is discussed.

## 1. Introduction

Ascorbic acid (or its salt form ascorbate) is an essential enzyme cofactor, participating in collagen and norepinephrine synthesis, as well as a potent antioxidant and free radical scavenger [1,2]. Despite its importance, anthropoid primates, teleost fish, Guinea pigs, some bats and Passeriformes bird species, lack a functional L-gulono-γ-lactone oxidase (GLO) gene and, thus, the ability to catalyse the last step of ascorbic acid synthesis [3,4]. The scattered loss of the GLO gene was suggested to evolve neutrally, with ascorbic acid requirements being met by dietary intake [3]. Unlike ascorbate, uric acid is largely regarded as end-product of purine metabolism and metabolic waste [5]. Yet, human-biased research has suggested alternative physiological functions for uric acid or urate (uric acid salts), such as antioxidant and free radical scavenger, similarly to ascorbic acid, counterbalancing disease-related oxidative stress [6-8].

Ascorbic and uric acid serve as substrates to active transporters that belong to the Nucleobase-Ascorbate Transporter (NAT) family; NAT family members, which include Solute carrier family 23 (Slc23) transporters, display a very broad evolutionary distribution, from bacteria to metazoans [1]. Sodium-dependent Slc23 transporters were suggested to maintain vertebrate ascorbic acid levels: with Slc23a1 responsible for renal and intestine ascorbic acid absorption across brush border epithelia and Slc23a2, which exhibits a widespread expression, responsible for ascorbic acid supply to target tissues and cells, including neurons [1,9,10]. In vertebrates, two additional transporters were identified, the orphan transporter Slc23a3 and the nucleobase transporter Slc23a4, described in rats but found pseudogenized in humans [1,11]. Rat Slc23a4 is restricted to the small intestine and was shown to predominantly mediate sodium-dependent uracil transport [11]. Slc23a3, on the other hand, is expressed in proximal renal tubules but is unresponsive to both ascorbic acid and nucleobases [1].

NAT family members display a highly conserved signature motif: [Q/E/P]-N-x-G-x-x-x-x-T-[R/K/G] [1,11-13]. Functional characterization of distinct member of the NAT family highlighted the importance of the first amino acid position of the motif in the definition of substrate specificity [1,11-13]. Thus, these transporters can be divided into three functional groups. Slc23a1, Slc23a2 and Slc23a3 belong to the ascorbate group and harbour a conserved proline (P) in the first amino acid position of the signature motif [1]. A glutamate (E) residue defines the uracil group, including bacterial anion symporters and Slc23a4, while a glutamine (Q) is present in the bacterial uric acid and/or xanthine anion symporters [1,11-13]. Topology predictions, using proteins from the distinct groups, suggest a cytosolic location of the motif [1, 11, 12]. Here we revise the portfolio of NAT family members in vertebrates.

## 2. Materials and Methods

### 2.1 Sequence mining

Slc23 protein sequences were obtained from the GenBank database by BLASTp using human Slc23a1 (NP_005838), Slc23a2 (NP_976072) and Slc23a3 (NP_001138362) sequences as query. The sequences from *Lethenteron japonica* (Japanese lamprey) were obtained from the Japanese Lamprey Genome Project (http://jlampreygenome.imcb.a-star.edu.sg/). The sequences from *Leucoraja erinacea* (little skate) and *Scyliorhinus canicula* (small-spotted catshark) were obtained as nucleotide sequences from SkateBase (http://skatebase.org) and translated. Accession numbers for all the retrieved sequences are listed in Supplementary Table 1 (Table S1).

### 2.2 Synteny

*Slc23* genes were mapped onto the respective species genomes, using the latest genome assemblies available in GenBank database (https://www.ncbi.nlm.nih.gov/). Genes flanking *Slc23a1, Slc23a2, Slc23a3, Slc23a4* and the novel *Slc23a*-like gene were identified and mapped using the human loci as reference. If the target gene was not found, the syntenic genomic region was retrieved using neighboring genes as reference.

### 2.3 Phylogenetic Analysis

The retrieved amino acid sequences were aligned using the MAFFT software web service (http://mafft.cbrc.jp/alignment/software/) in default automatic settings, with L-INS-I refinement method [14]. The alignment was stripped from columns with at least 20% of gaps resulting in an alignment with 75 sequences and 526 positions. The output alignment was used to construct a phylogenetic tree using PhyML 3.0 with Smart Model Selection web service (http://www.atgc-montpellier.fr/phyml-sms/) in default settings, which selected the LG +G+I+F model [15]. Bayesian-like transformation of aLRT was selected to assess branch support. An additional phylogenetic tree, with automatic bootstrapping (552 bootstraps) and using the default hill-climbing algorithm, was constructed with RAxML, available in CIPRES Science Gateway.

### 2.4 Membrane topology prediction

Slc23a5 membrane topology predictions were performed for selected species, using distinct prediction web servers: TMHMM Server v. 2.0 (http://www.cbs.dtu.dk/services/TMHMM/) and TMPred (https://embnet.vital-it.ch/software/TMPRED_form.html). Human Slc23a1 was used as control.

### 2.5 Filtering and quality control of Sequence Reading Archive (SRA) datasets

For RNA-seq analysis, we included 39 SRA datasets from brain, intestine, kidney, liver, and testis tissues of *Homo sapiens* (human), *Mus musculus* (mouse), *Gallus gallus* (chicken), *Anolis carolinensis* (American green anole), *Xenopus tropicalis* (western clawed frog), *Danio rerio* (zebrafish), *Lepisosteus oculatus* (spotted gar) and *Oryzias latipes* (Japanese medaka), which were obtained from the National Center for Biotechnology (NCBI) Sequence Read Archive (*SRA*) (https://www.ncbi.nlm.nih.gov/sra/) (Table S2). The read quality assessment of each SRA dataset was performed using FastQC software (https://www.bioinformatics.Babraham.ac.uk/projects/fastqc/). Trimmomatic 0.36 [17] was used to trim and drop reads with quality scores below 5, at leading and trailing ends, with an average quality score below 20 in a 4 bp sliding window and with less than 36 bases length.

### 2.6 Relative gene expression levels

To assess the relative gene expression of *slc23* genes from the selected species, reference sequences and the corresponding annotations were collected from NCBI and Ensembl [18] databases. For human, mouse, lizard, zebrafish and spotted gar, the GTF and DNA files (genome) were retrieved from Ensembl (Release 89) [18]. For chicken and western clawed frog, transcript sequences (coding and non-coding) and mapping (gene/transcript) from NCBI Refseq database were used after filtering [19] (for each species the rna.fasta file from ftp://ftp.ncbi.nih.gov/genomes/refseq/ was used – See Table S3 and S4 for detail). The SRA reads of each specimen were aligned to the reference using Bowtie2 (default parameters)[20], and gene expression was quantified as transcript per million (TPM) using RSEM v.1.2.31 (RNA-seq by Expectation Maximization) software package [21]. Isoform expression levels for each gene were summed to derive the TPM values. For genes with low relative expression values, TPM values were log2-transformed after adding a value of one; TPM values < 0.5 were considered unreliable and substituted with zero.

### 2.7 PCR-based validation of Slc23a5 expression

Total RNA extraction from *X. tropicalis* and *D. rerio* tissues was performed using the Illustra RNAspin Mini RNA Isolation Kit animal tissues protocol with on-column DNAseI digestion (GE Healthcare). RNA quality was visually assessed by electrophoresis and sample concentration was determined using the Take3 micro-volume plate system (BioTek). First-strand cDNA was synthesized from 250ng of total RNA using the iScriptTM cDNA Synthesis Kit (Bio-Rad), according to the manufacturer’s instructions. Gene expression of *Slc23a5* was analyzed by PCR using the intro flanking expression primer pairs 5’-AGACGTTGAAGCCGTTACTGA-3’ and 5’-TGCTATTATGCCTCCAAAGGCT-3’ for *X. tropicalis* and 5’-GGTGGAATGTTCCTCGTCAT-3’ and 5’-GATGAACATGTGGGTGGTGA-3’ for *D. rerio*. PCR reactions were carried out on a tissue panel containing heart, skin, stomach, intestine, liver, kidney, ovary and testis tissues for *X. tropicalis* and brain, heart, skin, gut, liver, kidney, ovary and testis tissues for *D. rerio*. The PCR was performed with Phusion^®^ Flash high fidelity Master Mix (Thermofisher), 1μM of each primer and 25ng/μl of template cDNA, with the following parameters: initial denaturation at 98°C for 1s, annealing at 58°C for *D. rerio* or 66°C for *X. tropicalis* primers and elongation at 72°C for 4s, with 30 cycles.

## 3. Results

### 3.1 Identification of a novel member of the NAT family in neopterygians and amphibians

Through database mining and phylogenetic analysis, we detected the presence of a novel gene, *slc23a5*, found in neopterygians (Holostei and Teleostei) and amphibians (Figure 1 and Figure S1). No *slc23a5* sequence was retrieved from chondrichthyans, chondrosteans (basal actinopterygians) nor from the sarcopterygian *Latimeria chalumnae* (coelacanth). Subsequent loss of *slc23a5* in chondrichthyans was further supported by the conservation of the genomic locus in *Callorhynchus milii* (Australian ghostshark) (Figure S2). Regarding coelacanth, two scaffolds, denoting conservation of the flanking genes, were found; while, no information was retrieved regarding the chondrostean loci (Figure S2). The remaining genes of the *slc23* family were retrieved and genomic loci were found generally conserved across vertebrate species, with exception of the *slc23a4* loci in fish (Figures S3-S6). The branching pattern of our phylogenetic analysis suggests that after the expansion of the *slc23a* gene family *slc23a5* was subsequently lost in sharks, basal actinopterygians, coelacanth and amniotes (Figure 1 and Figure S2). In agreement, we obtained three phylogenetic clusters possibly corresponding to three ancestral lineages (Figure 1 and Figure S1): one composed of typical *slc23a1* and *slc23a2* genes and a second group including the previously reported *slc23a4* and the novel *slc23a5* genes, both outgrouped by invertebrate sequences. A third cluster, which outgroups the remaining *slc23* genes, included the highly divergent *slc23a3* genes, mirrored by the long branch lengths (Figure 1).

**Figure 1.**
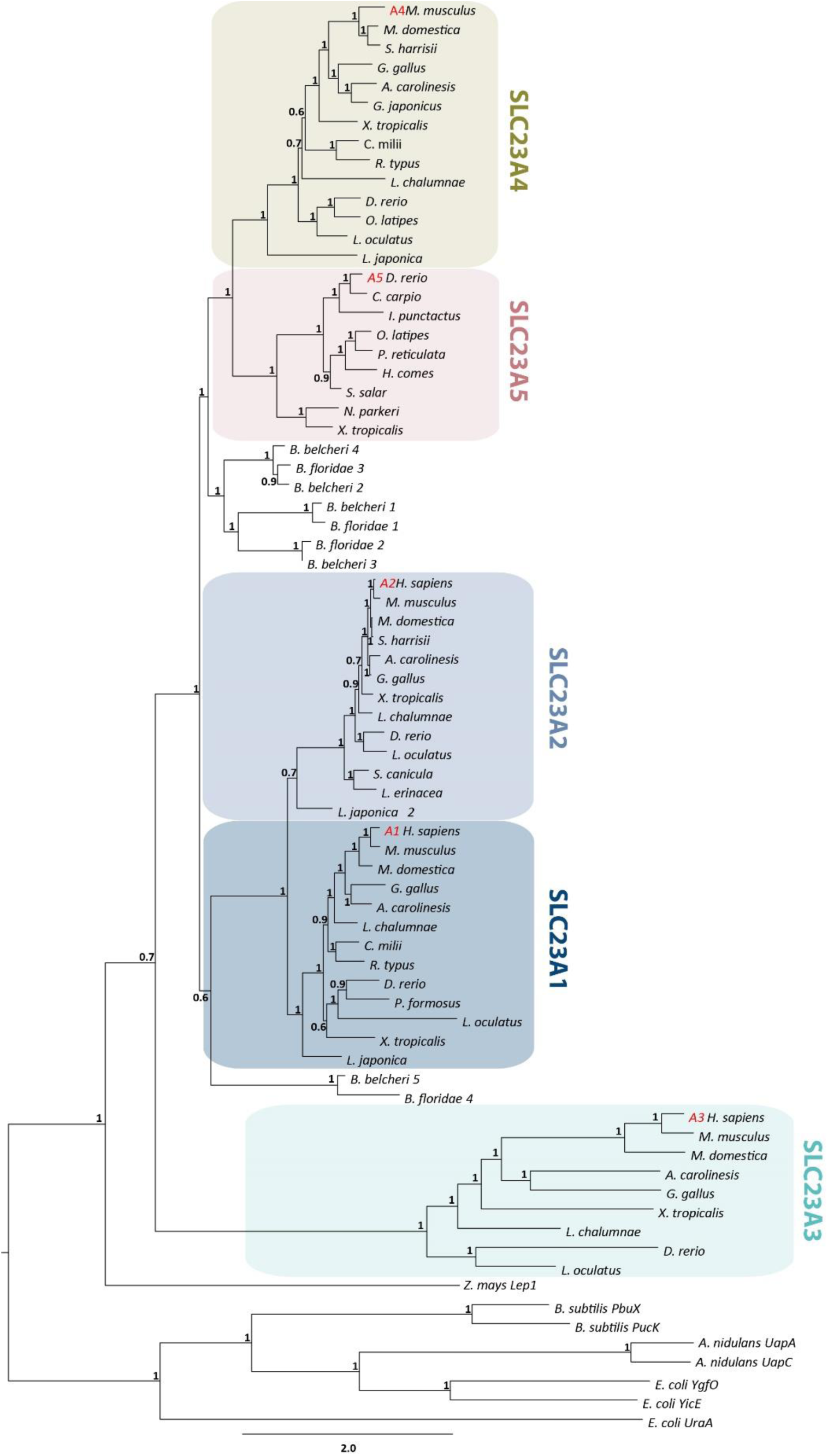
Maximum likelihood phylogenetic tree describing relationships among Nucleobase-Ascorbate Transporter (NAT) family members. Node values represent branch support using the aBayes algorithm. Accession numbers for all sequences are provided in Supplementary Materials.

### 3.2 Slc23a5 signature motifs

Further comparative protein sequence analysis highlighted the presence of a NAT family signature motif with a conserved Q in the substrate determining position. According to previous functional characterization of NAT family members, this residue is found in uric acid and/or xanthine transporters, thus placing Slc23a5 in the uric acid and/or xanthine group (Figure 2)[1,11-13]. Besides the typical signature motif, a carboxyl terminus tripeptide motif was identified in some SLC23 family members: a S/T-X-Ø tripeptide. S/T-X-Ø motifs were shown to interact PDZ domains of scaffolding proteins belonging to the Sodium/Hydrogen Exchanger Regulatory Factor, NHERF, family [22]. S/T-X-Ø are found highly conserved across species in Slc23a1 and Slc23a4, in agreement with their renal and intestinal expression and role in ascorbic acid and nucleobase absorption (Figure S7). Slc23a5, on the other hand, exhibits a highly conserved tripeptide motif in Otocephala (Cypriniformes, Characiformes and Siluriformes) but not Osteoglossomorpha (Figure 3A). The Euteleostei Esociformes and Salmoniformes exhibit an inverted motif while in the Neoteleostei no conservation was found. Finally, topology predictions yielded similar transmembrane profiles for Slc23a5 sequences from *D. rerio, O. latipes, Salmo salar* (Atlantic salmon) and *X. tropicalis*, when compared to human Slc23a1 (Figure S8). The NAT motif was predicted cytosolic in all analyzed sequences. This suggests a conserved cellular import topology, as observed in other family members.

**Figure 2.**
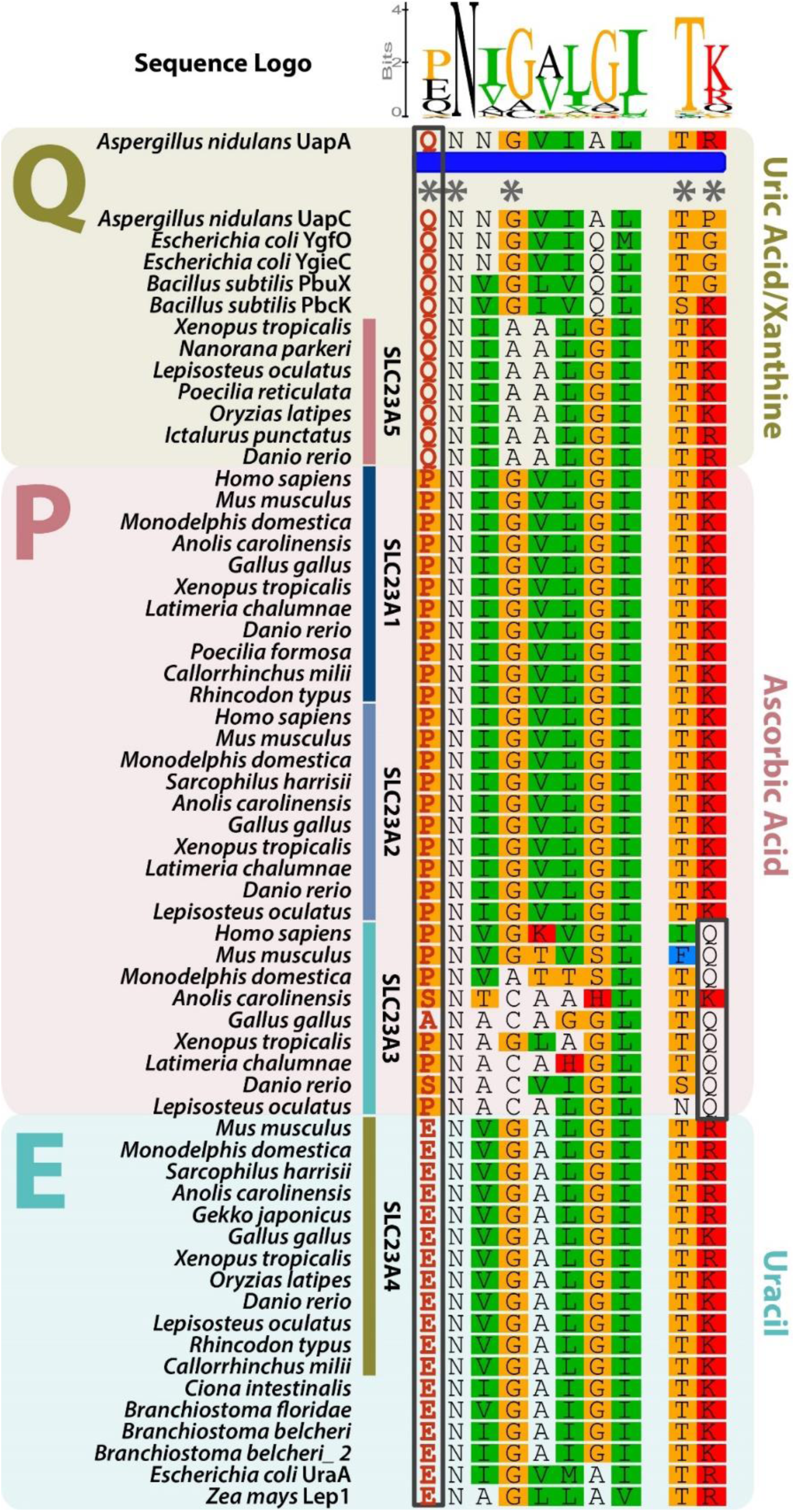
Alignment of the signature motif of the Nucleobase-Ascorbate Transporter (NAT) family. NAT family members display a highly conserved signature motif defining substrate specificity: [Q/E/P]-N-x-G-x-x-x-x-T-[R/K/G]. Asterisks indicate conserved positions. The first amino acid position, with residues highlighted in red within a black box, defines substrate preference: Q – Uric acid/xanthine group; P – Ascorbate group and E – Uracil group. The mirror Q in the Slc23a3 group is also highlighted by a black box.

**Figure 3.**
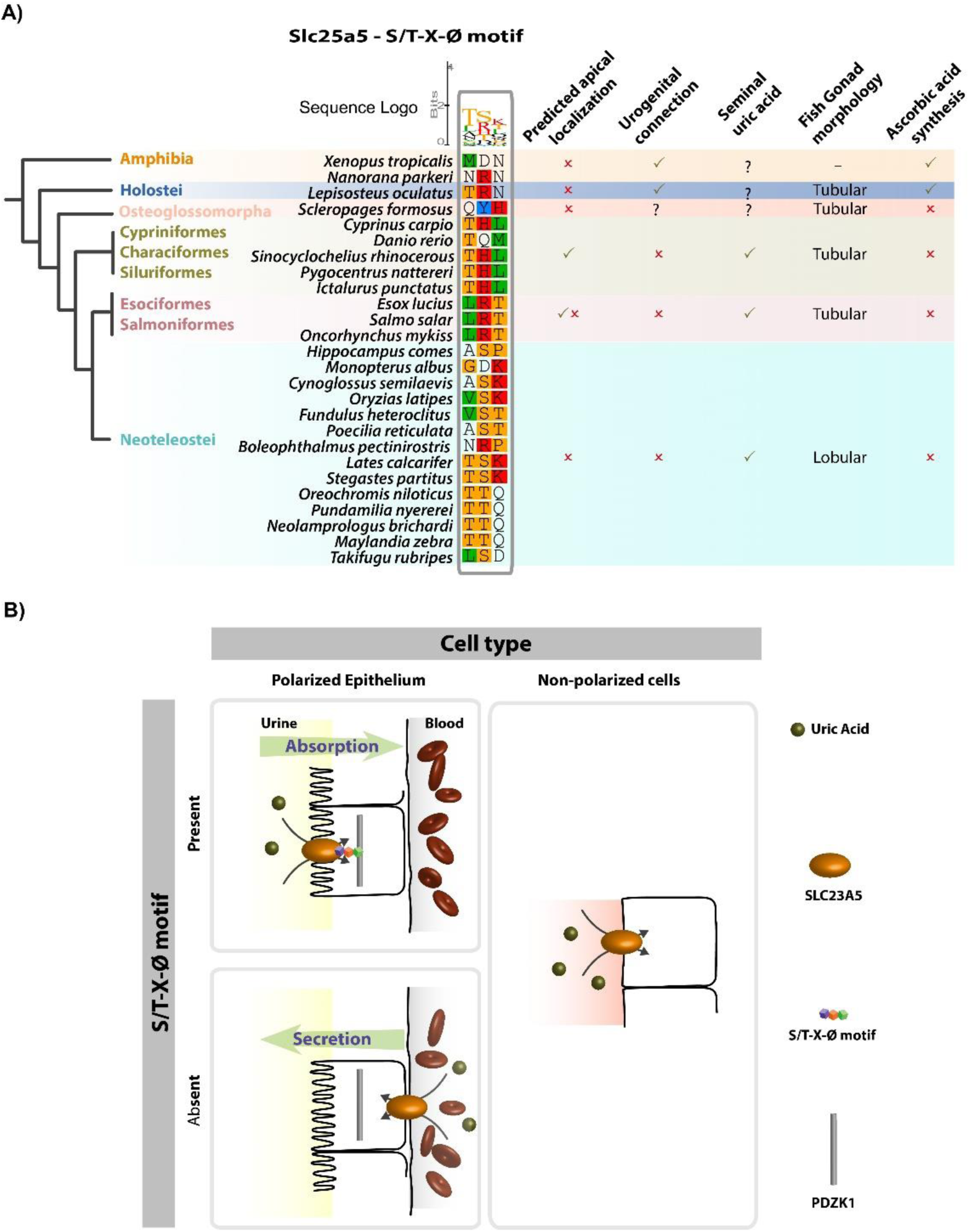
The S/T-X-Ø tripeptide provides insight into the cellular function of Slc23a5 homologues from fish and amphibians. (A) Alignment of the PDZ-binding tripeptide in Slc23a5 homologues from fish and amphibians. The S/T-X-Ø tripeptide is found in Cypriniformes, Characiformes and Siluriformes while the Euteleostei Esociformes and Salmoniformes exhibit an inverted motif. Predicted cellular localization is indicated as well as morphological and physiological adaptations among the different fish groups. (B) Schematic representation of tripeptide-dependent uric acid flows in polarized epithelia and non-polarized cells. In polarized epithelia (i.e. kidney) the presence of the tripeptide motif suggests interaction with the scaffolding protein PDZK1 which localizes apically, in the absorptive side.

### 3.3 Slc23a5 expression patterns

To gain further insight into the functional relevance of this additional gene, we collected RNA-seq data and determined relative expression levels of *Slc23* genes. Gene expression analysis suggested that, in *D. rerio* and *X. tropicalis*, slc23a5 is mostly expressed in kidney and testis, further confirmed by PCR (Figure 4 and Table S5). Expression in brain was also detected in *D. rerio* and was found to be the exclusive expression site in the Neoteleostei *O. latipes*. The holostean *L. oculatus*, on the other hand, yielded no significant expression in the tested tissues.

**Figure 4.**
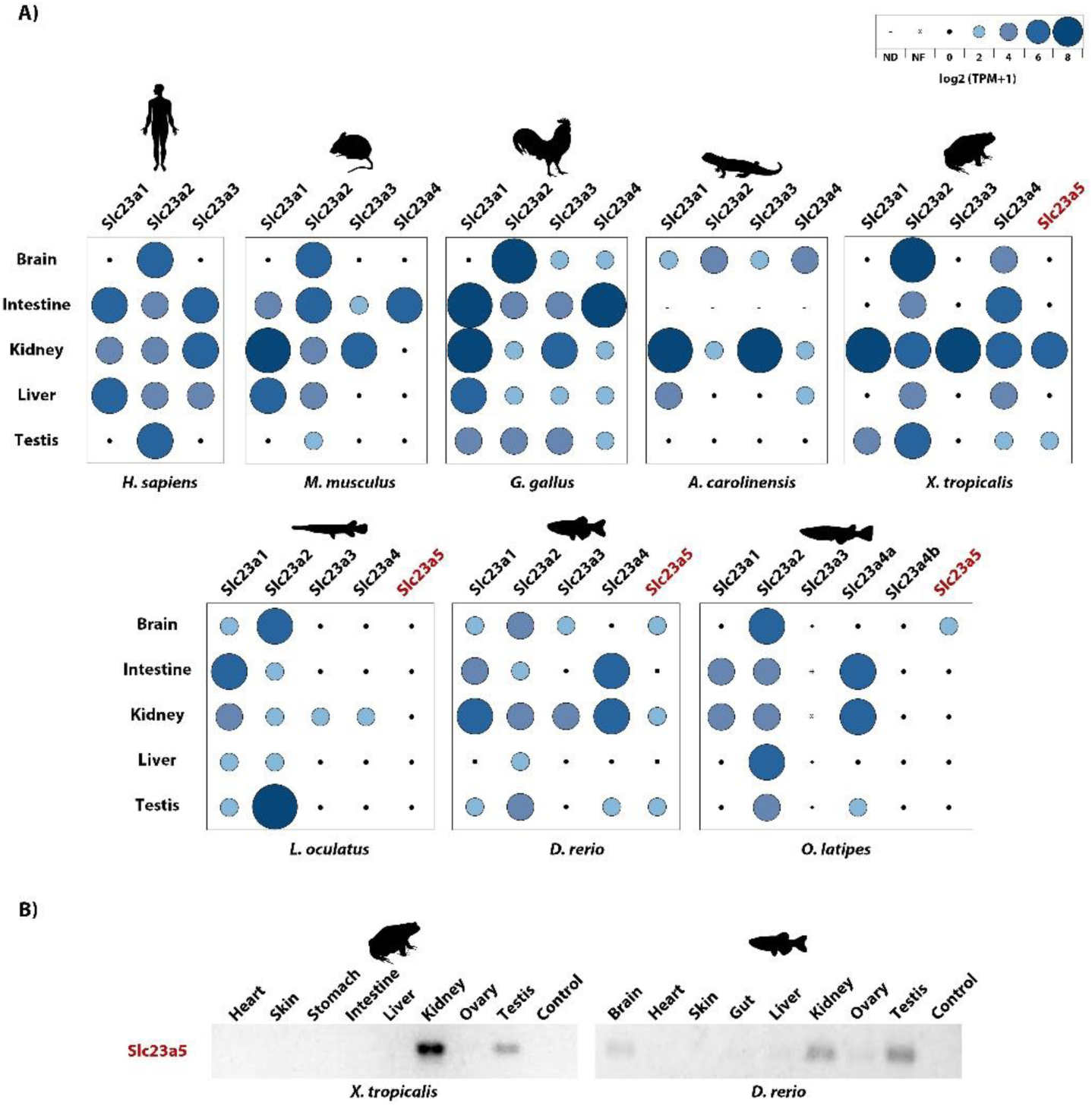
Expression patterns of *Slc23* isoforms across vertebrates. (A) RNAseq-based quantification. Relative gene expression levels are provided as log_2_-transformed transcript per million (TPM) values, from 0 to 8 TPM’s, the NF and ND means gene not found and no SRA data available to a specific tissue. (B) PCR-based amplification of *Slc23a5* transcripts in *D. rerio* and *X. tropicalis* tissues.

## 4. Discussion

Our findings indicate a restricted distribution of *slc23a5* to taxa that share an aquatic nature of their habitat. In fact, *slc23a5* was found to be expressed in kidney and testis of both *X. tropicalis* and *D. rerio*. In the latter, as well as in the Neoteleostei *O. latipes*, brain expression was also detected. Comparative analysis using previously characterized NAT family transporters placed this novel transporter in the uric acid and/or xanthine transporter group. In fact, similarly to uric acid and/or xanthine transporters, Slc23a5 proteins harbor a Q in the first position of the NAT family signature motif ([Q/E/P]-N-x-G-x-x-x-x-T-[R/K/G]). Yet, further functional studies are required to fully clarify the molecular function of Slc23a5 proteins, notably its uric acid and/or xanthine binding abilities.

Given the nature of NAT family transporter ligands, including ascorbic and uric acids, xanthine or nucleobases, renal Slc23a5 could participate in the homeostatic maintenance of excretion products, in osmotic regulation and/or in antioxidant processes. In fact, kidneys play a crucial role in the maintenance of both ascorbic and uric acid levels through the excretion and reabsorptions processes [6,22]. In kidney absorptive epithelium, transporters located on the filtrate-epithelial interface (apical side) interact with the scaffolding protein PDZ-Containing Kidney Protein 1 (PDZK1) [22]. PDZK1 belongs to the NHERF family of proteins which are mostly expressed in the apical side of absorptive epithelia of kidney, intestine and liver, and establish protein-protein interaction networks with receptors, transporters and channels [23,24]. Besides its scaffolding role, NHERF family members were also shown to modulate the activity of protein assemblages [24-26]. Within Slc23 family members, S/T-X-Ø motifs are found in Slc23a1 and Slc23a4 proteins, which are present in renal and intestinal polarized epithelia and participate in ascorbic acid and nucleobase absorption, and absent in Slc23a2. Curiously, a highly conserved PDZK1 interacting motif (S/T-X-Ø) was also detected in Slc23a5 from Cypriniformes, Characiformes and Siluriformes, but not in other fish groups or amphibians. No relationship between salt and freshwater environments was found. Thus, the presence/absence of this peptide signal might reflect distinct flows in the kidney: an apically expressed Slc23a5 likely participates in absorptive flows, while a basolateral expression would be related to secretion (Figure 3B).

In the human proximal renal tubules, several transporters, localized to specific cellular regions (apical or basolateral), seem to cooperate for efficient urate handling: named the urate transportome [6,22]. Thus, if the predicted uric acid transport activity is confirmed, *X. tropicalis* and *D. rerio* Slc23a5 could illustrate the two opposing mechanisms: while *D. rerio* captures, *X. tropicalis* secretes uric acid into renal filtrate. In agreement, *X. tropicalis, D. rerio*, as well as *L. oculatus*, appear to lack the full complement of renal urate transporters (Table S6). A similar scenario is observed for birds and reptiles; however, their excretory system represents a particular adaptation for uric acid excretion [27,28]. Besides simple excretion, and given that high concentrations of uric acid were detected, and suggested to act as the main non-enzymatic antioxidant mechanism, in seminal fluid of several fish species (including Cypriniformes, Esociformes, Salmoniformes and the Neoteleostei Gadiformes and Perciformes [29-32]), this putative uric acid transport activity could be maintained for reproductive purposes, in agreement with the observed testicular expression. In fact, in external fertilizers, such as most fish and amphibians, reactive oxygen species were suggested to decrease motility and thus fertilisation success [29]. Yet, regarding amphibians, no information on seminal fluid composition was retrieved [33]. Nonetheless, in vitro assays suggested that other antioxidant species, such as ascorbic acid or vitamin E, were unsuitable at offsetting oxidative stress and improving sperm motility in *Litoria booroolongensis* (Booroolong frog) and suggested uric acid as a pertinent candidate [33]. In this context, the absence of *slc23a5* sequences in Chondrichthyes could be due to their internal fertilization strategies derived from ovoviviparous or viviparous reproductive modes [34]. Accordingly, viviparity is also observed in the sarcopterygian *L. chalumnae* [34].

Besides reproductive categories, other morphological and metabolic traits could justify the seemingly scattered retention of *slc23a5* among fish in the context of the proposed uric acid transport activity; for instance, specific anatomical features such as the presence of a urogenital connection: described in Elasmobranchii and in most non-teleost Osteichthyes, yet absent in Cyclostomata and Teleostei [35]. By connecting testicular and renal ducts, antioxidant requirements could be met by renal secretion into urine. In agreement, in the Chondrostei *Acipenser ruthenus*, full sperm maturation and motility was shown to require dilution by urine [36]. An urogenital connection is also present in amphibians [35]. Additionally, other antioxidant mechanisms could exist. Besides uric acid, ascorbic acid is also present in seminal fluid of *Oncorhynchus mykiss* (rainbow trout) [37]. Seminal ascorbate was reported to be affected by seasonal variations and semen quality was shown to improve upon dietary ascorbic acid supplementation [37,38], contrasting with the amphibian *L. booroolongensis* [33]. In the case of non-teleost fish, such as Chondrostei (*A. ruthenus*) and Holostei (*L. oculatus*), a functional GLO gene is retained, providing endogenously synthesised ascorbate [39]. Curiously, no gonadal expression was detected in the Neoteleostei *O. latipes*. Neoteleosts exhibit varied reproduction modes ranging from external fertilizers to internal fertilizers with viviparity [34,40-42]. Also, neoteleosts were shown to exhibit a distinctive gonadal morphology, the lobular testis, whereas the remaining bony fish exhibit tubular testis [43]. Although still obscure, the absence of testicular slc23a5 expression in the externally fertilizer *O. latipes* could be due to specific morphological adaptations of neoteleosts.

A similar role could be postulated for neuronal uric acid [44]. Active swimming behaviour requires a high metabolic rate, mirrored by an increased oxygen consumption, subsequently leading to reactive oxygen species production [45]. In agreement, fish antioxidant levels were shown to be correlated with their swimming activity [45]. Neuronal cell populations are particularly sensitive to oxidative stress, thus, uric acid could participate in the neutralization of oxidative species to by-pass oxidative stress, notably in GLO-deficient fish. Yet, information on neuronal uric acid levels in fish is currently absent.

## 5. Conclusion

Taken together, these observations put forward the retention of an additional *Slc23* gene, in neopterygians and amphibians, with restricted expression patterns and presumed uric acid transport function. Yet, extensive comparative functional analysis should be conducted to clarify the physiological relevance of this novel receptor in aquatic organisms.

## Supplementary material

Supplementary tables and figures including synteny, phylogeny and topology analysis, as well as reference tables.

## Acknowledgments

This work was supported by Coral—Sustainable Ocean Exploitation (Norte-01-0145-FEDER-000036), a project by the North Portugal Regional Operational Program (NORTE 2020), under the PORTUGAL 2020 Partnership Agreement, through the European Regional Development Fund (ERDF) and by the European Regional Development Fund (ERDF) through the COMPETE - Operational Competitiveness Programme and POPH - Operational Human Potential Programme.

## A taxonomically-restricted Uric Acid transporter provides insight into antioxidant equilibrium

### Supplementary Figures

**Figure S1.**
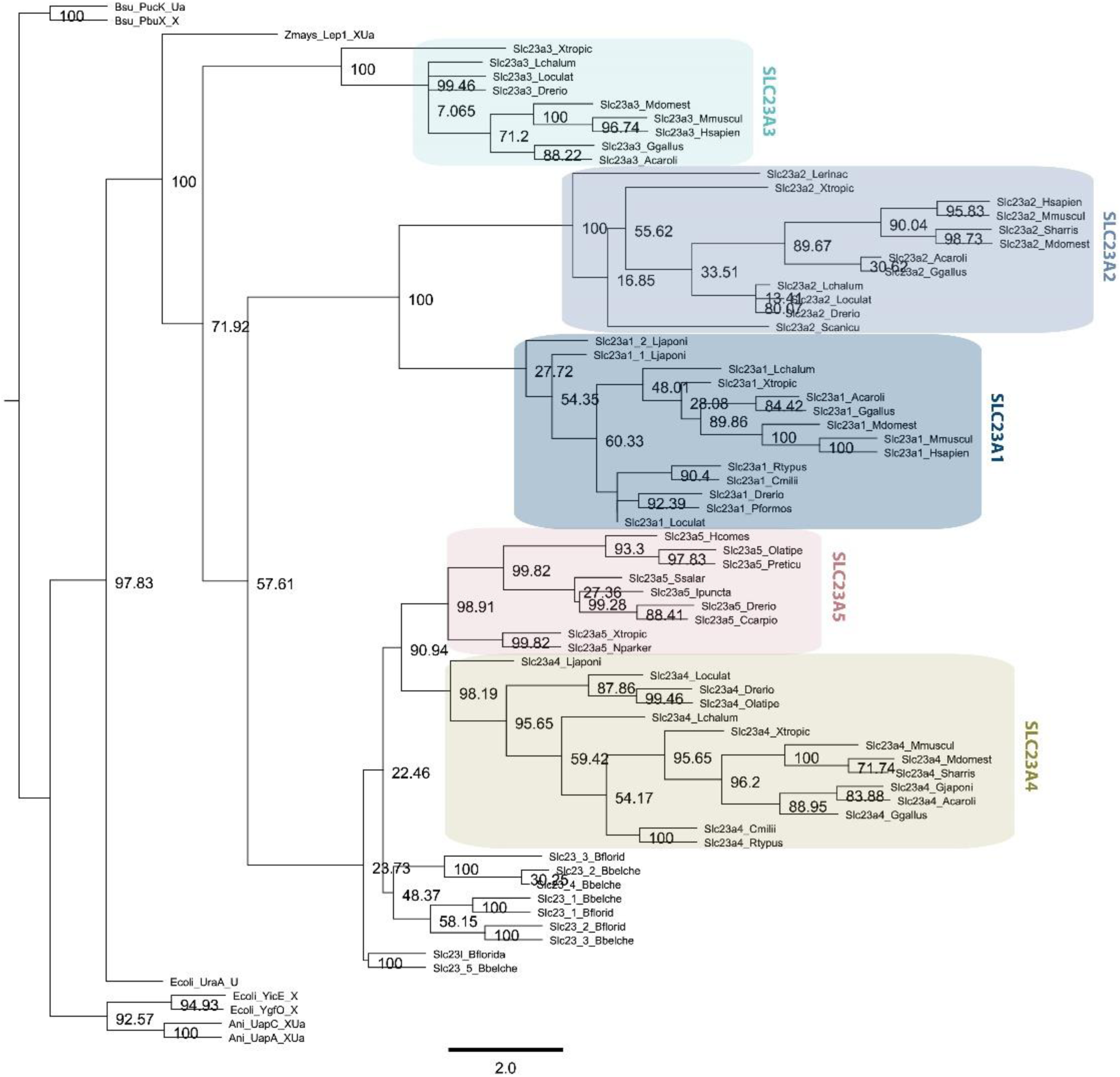
RaxML phylogenetic tree. Node values represent bootstrap values.

**Figure S2.**
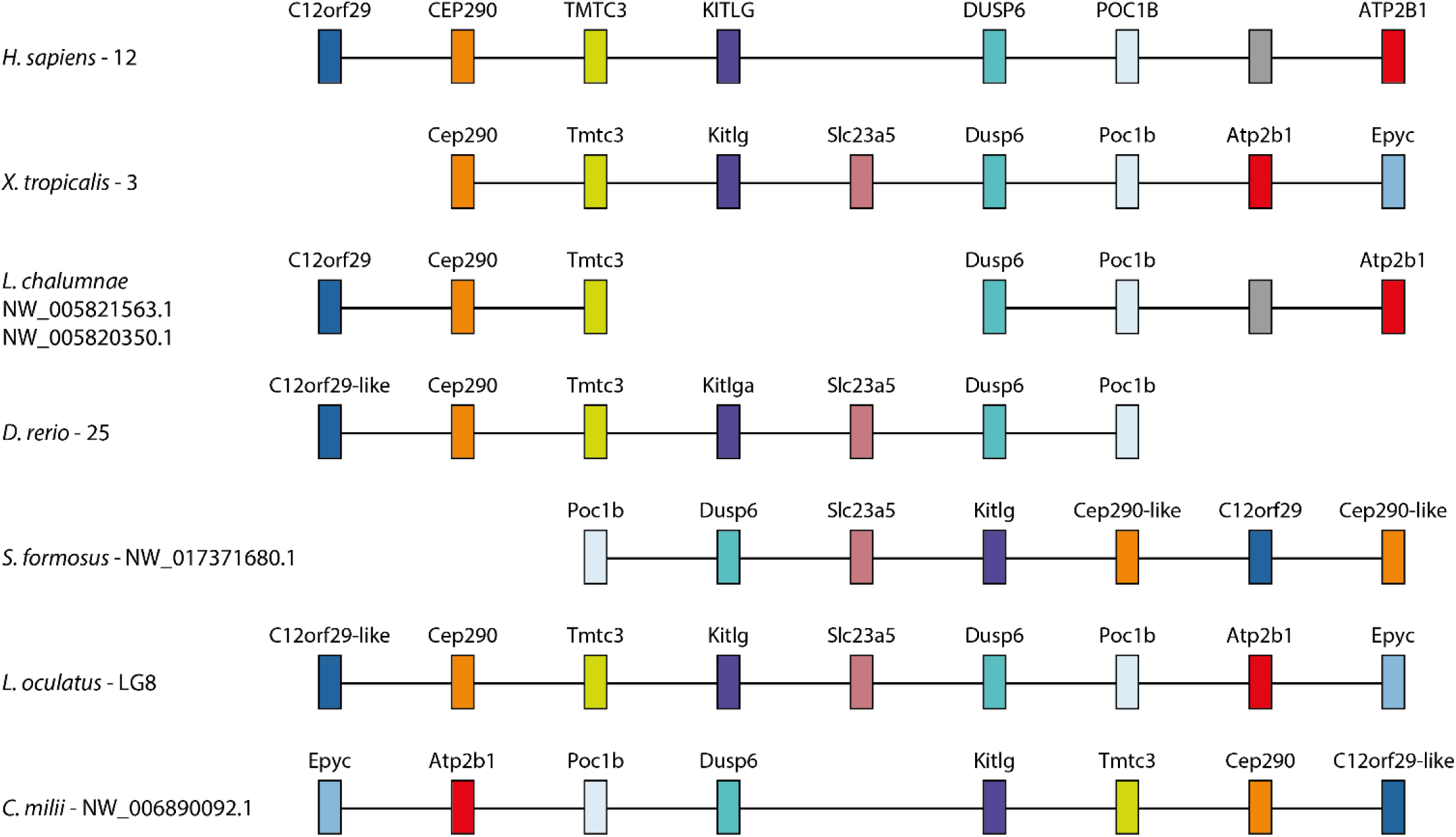
Synteny maps of *Slc23a5*. Conservation of the *Slc23a5* locus in major fish lineages, *Xenopus tropicalis* and *Homo sapiens*. Numbering after species corresponds to chromosome or scaffold. Information is presently absent for the Chondrostei *Acipenser ruthenus*. Grey boxes indicate non-conserved genes.

**Figure S3.**
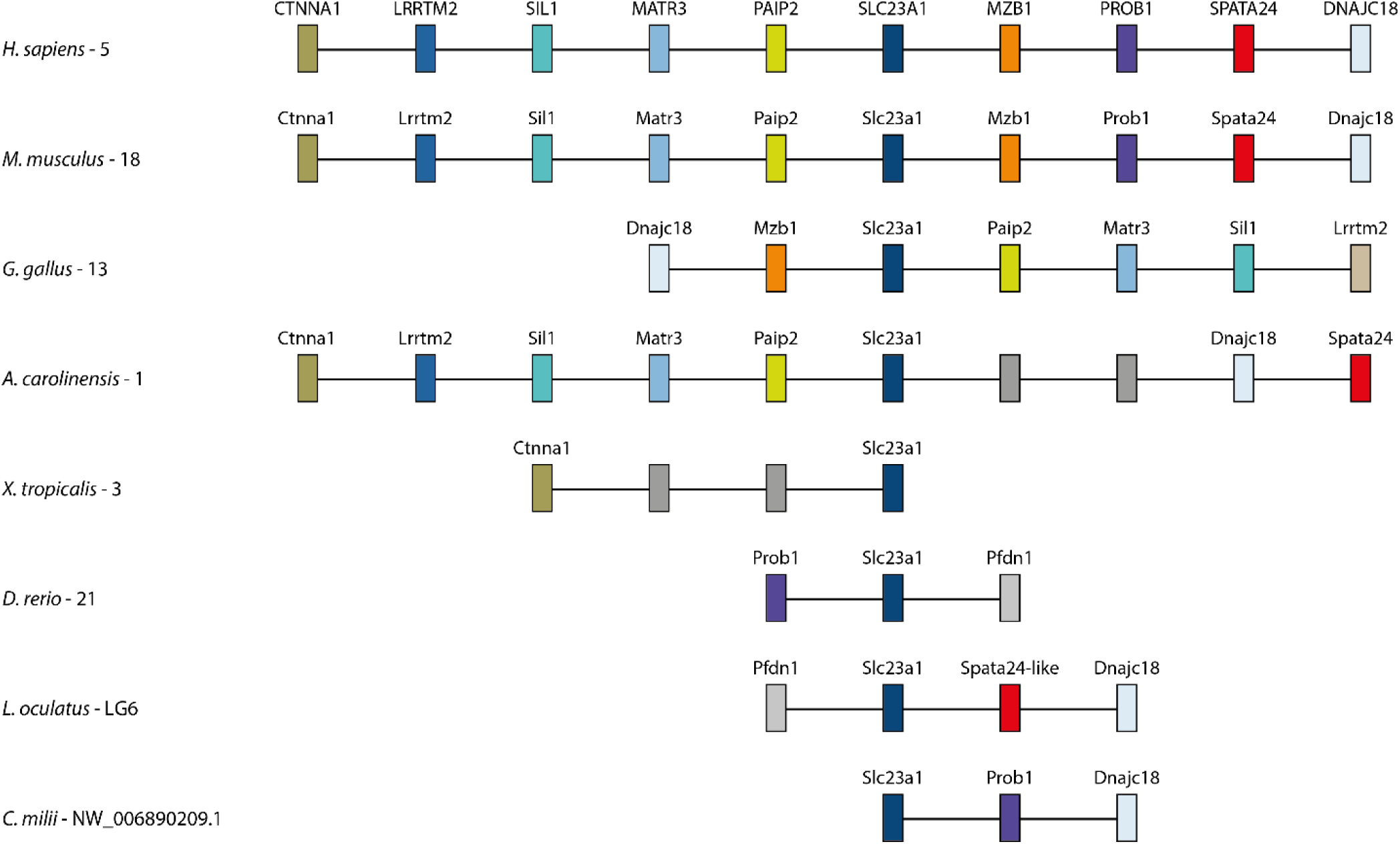
Synteny maps of *Slc23a1*. Conservation of the *Slc23a1* locus in major vertebrate lineages. Numbering after species corresponds to chromosome or scaffold. Grey boxes indicate non-conserved genes.

**Figure S4.**
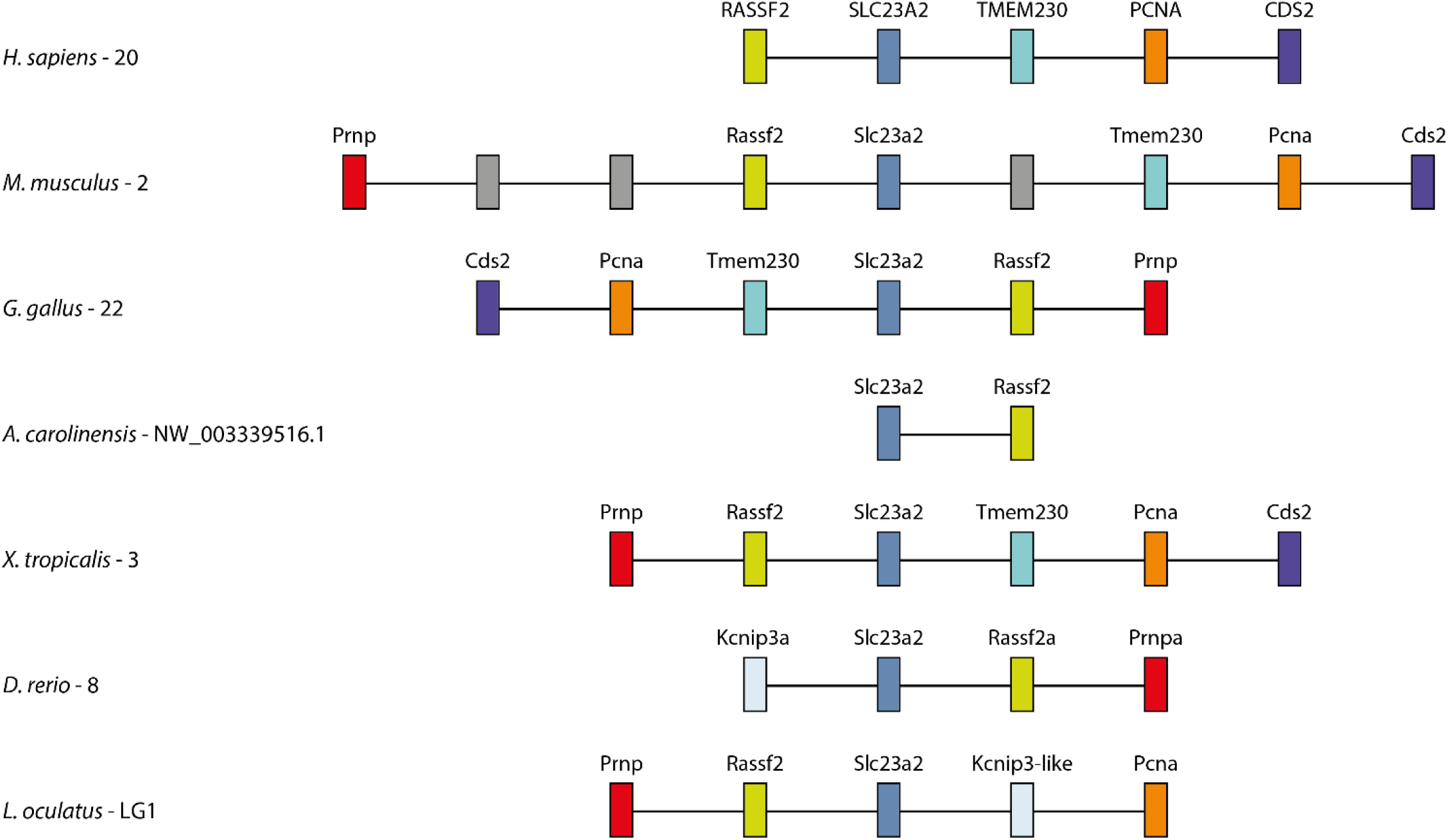
Synteny maps of *Slc23a2*. Conservation of the *Slc23a2* locus in major vertebrate lineages. Numbering after species corresponds to chromosome or scaffold. Grey boxes indicate non-conserved genes.

**Figure S5.**
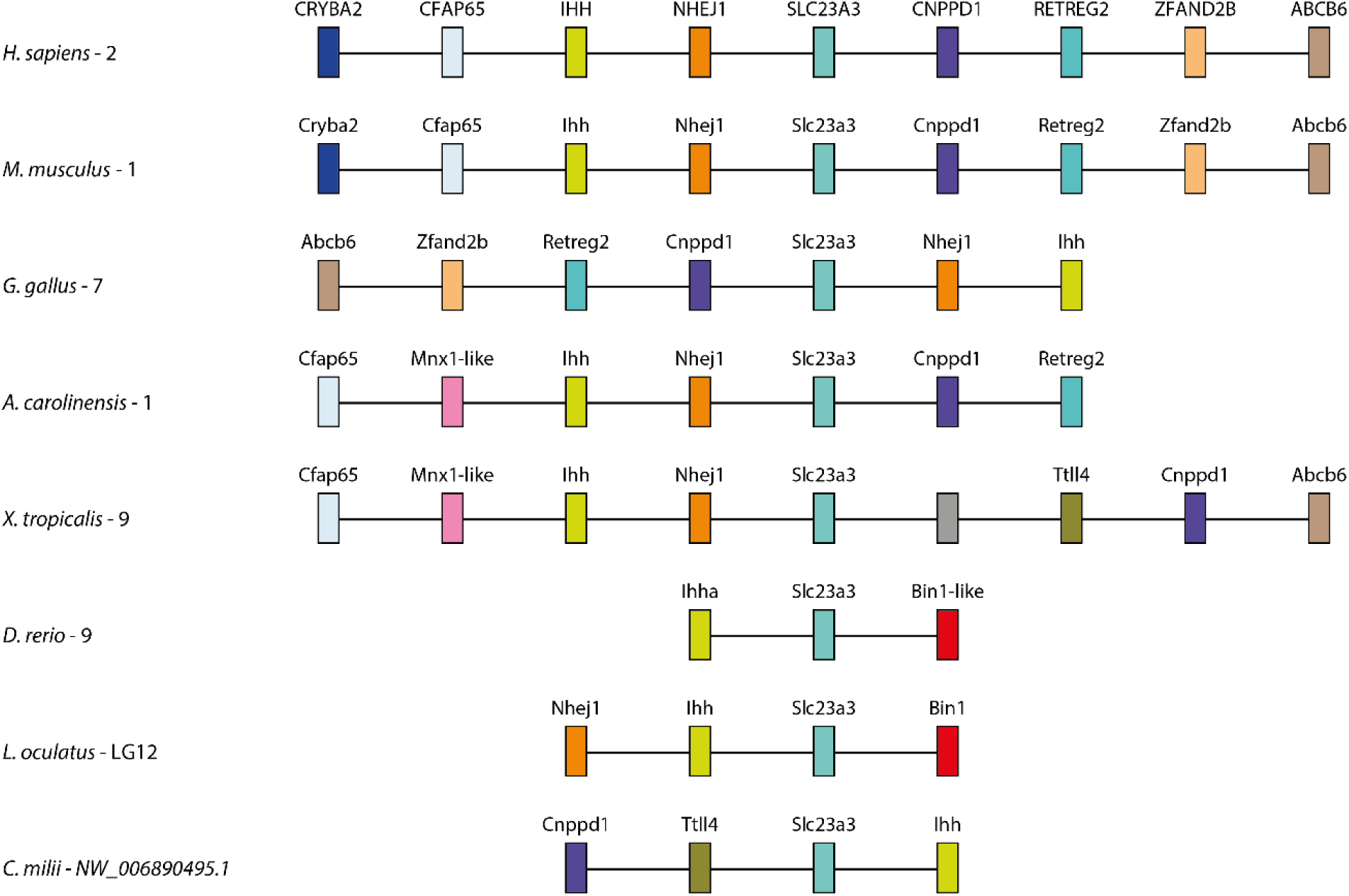
Synteny maps of *Slc23a3*. Conservation of the *Slc23a3* locus in major vertebrate lineages. Numbering after species corresponds to chromosome or scaffold. Grey boxes indicate non-conserved genes.

**Figure S6.**
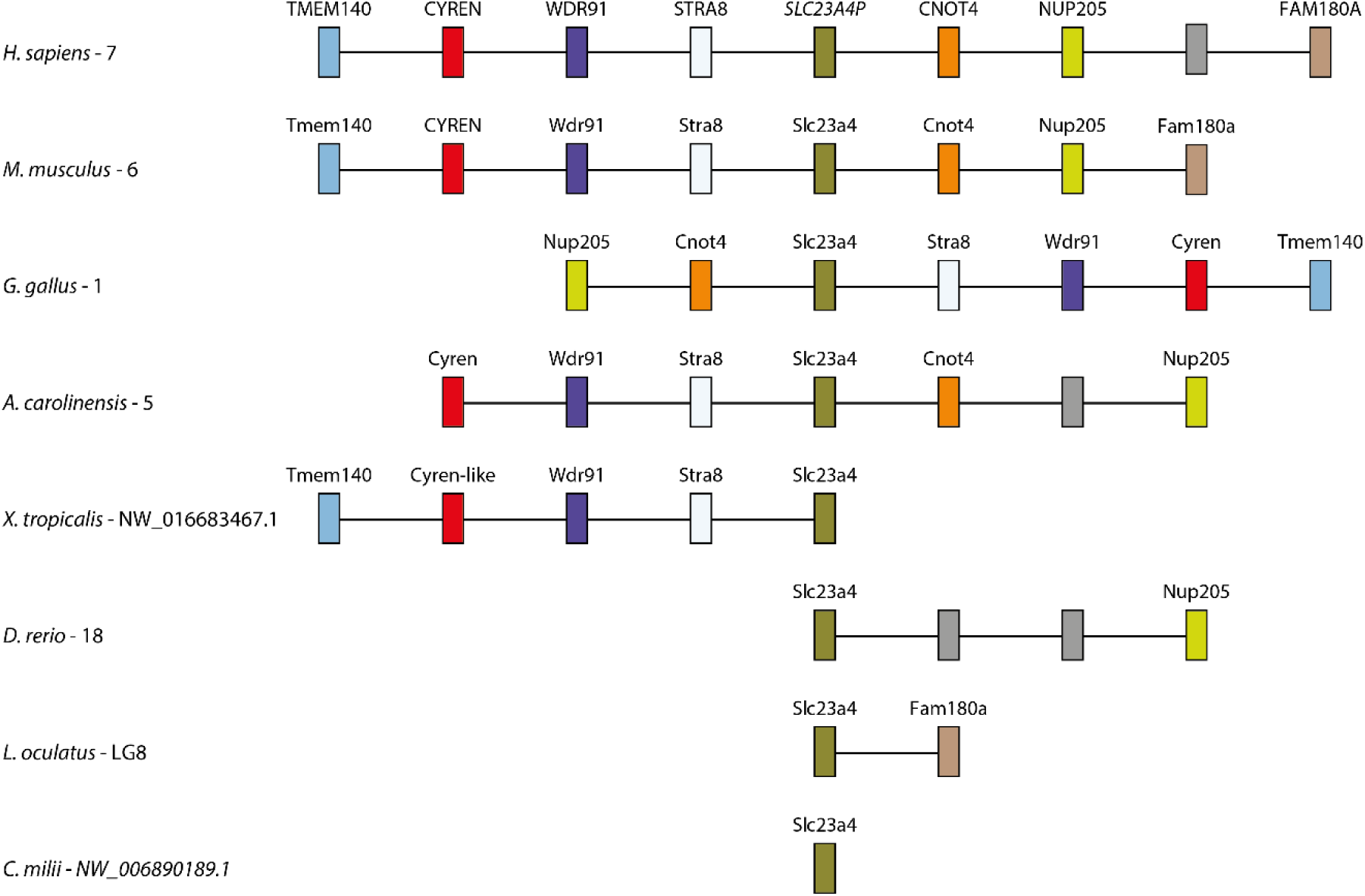
Synteny maps of *Slc23a4*. Conservation of the *Slc23a4* locus in major vertebrate lineages. Numbering after species corresponds to chromosome or scaffold. The human homologue is indicated as pseudogenized (*SLC23A4P*). Grey boxes indicate non-conserved genes.

**Figure S7.**
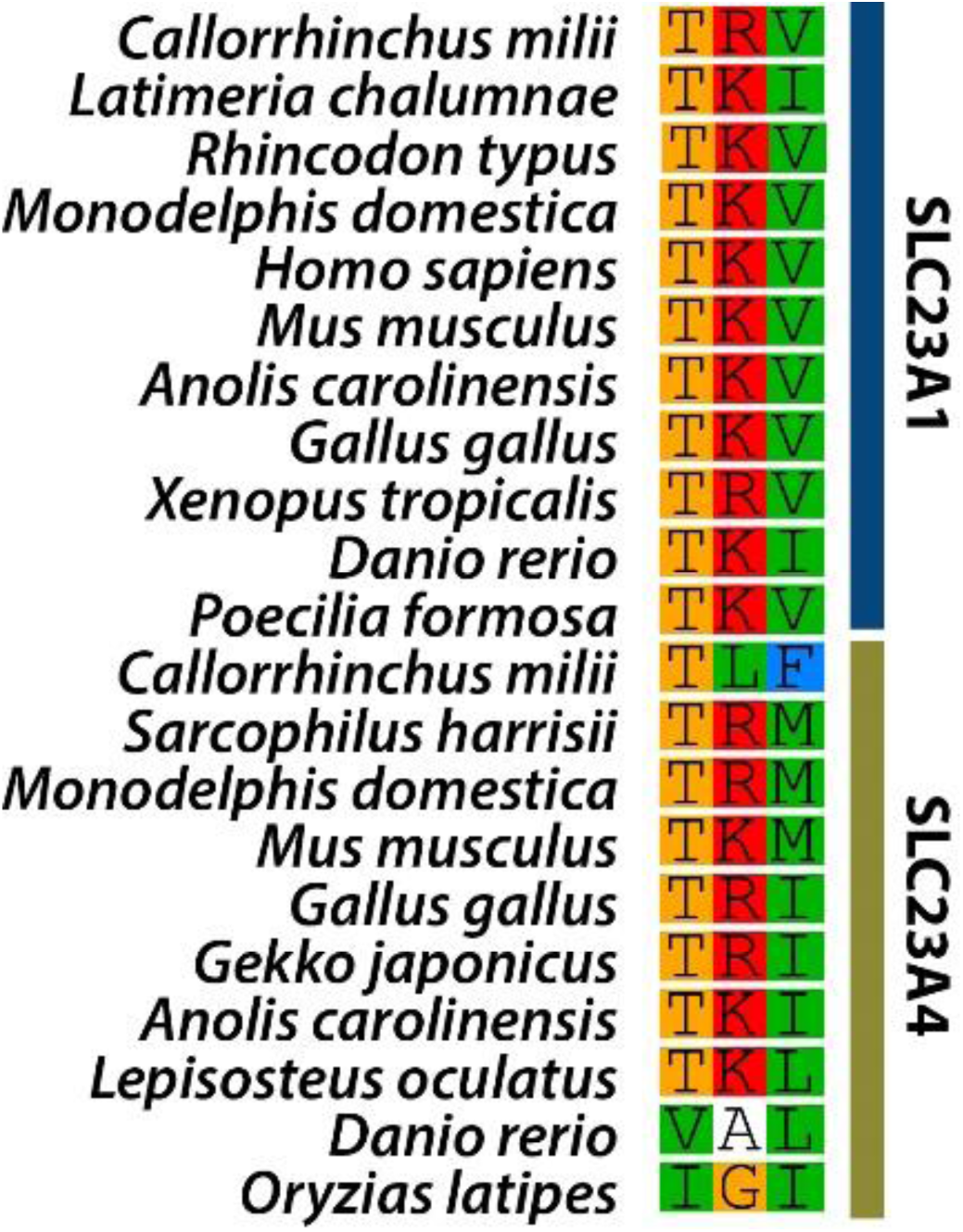
S/T-X-Ø tripeptide conservation in Slc23a1 and Slc23a4 sequences.

**Figure S8.**
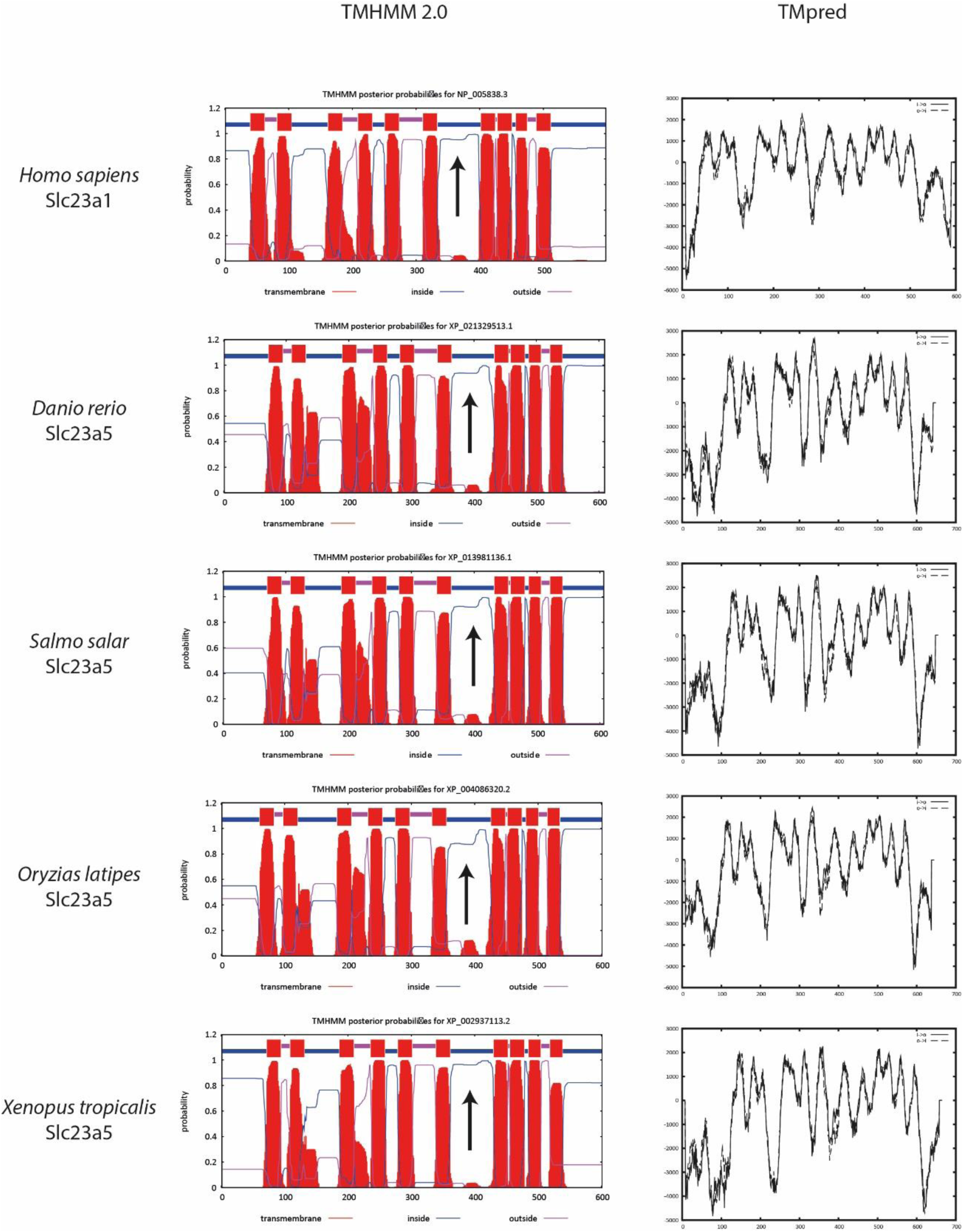
Topology prediction of Slc23a5 sequences from *D. rerio, O. latipes, Salmo salar* (Atlantic salmon) and *X. tropicalis*, and human Slc23a1 used as control. TMHMM Server v. 2.0 (left) and hydropathy plots (right) are provided. Arrows detone the position of NAT family motif.

### Supplementary Tables

**Table S1.** Accession numbers.

| Species | Gene | Accession Number | Common name |
| --- | --- | --- | --- |
| *Aspergillus nidulans* | UapA | CAA50681 | - |
| *Aspergillus nidulans* | UapC | CAA56190 | - |
| *Bacillus subtilis* | PbuX | KIX81273 | - |
| *Bacillus subtilis* | PucK | KIX81727 | - |
| *Escherichia coli* | YgfO | CDU41213.1 | - |
| *Escherichia coli O25b:H4* | UraA | ANK02789 | - |
| *Escherichia coli O25b:H4* | YicE | ANK04183 | - |
| *Zea mays* | Lep1 | AAB17501 | Maize |
| *Branchiostoma belcheri* | Slc23-like | XP_019630437 | Amphioxus |
| *Branchiostoma belcheri* | Slc23-like | XP_019636191 | Amphioxus |
| *Branchiostoma belcheri* | Slc23-like | XP_019633776 | Amphioxus |
| *Branchiostoma belcheri* | Slc23-like | XP_019616280 | Amphioxus |
| *Branchiostoma belcheri* | Slc23-like | XP_019646793 | Amphioxus |
| *Branchiostoma floridae* | Slc23-like | XP_002600710 | Florida lancelet |
| *Branchiostoma floridae* | Slc23-like | XP_002597306 | Florida lancelet |
| *Branchiostoma floridae* | Slc23-like | XP_002595086 | Florida lancelet |
| *Branchiostoma floridae* | Slc23-like | XP_002611161 | Florida lancelet |
| *Anolis carolinensis* | Slc23a1 | XP_008113637 | Green anole |
| *Callorhynchus milii* | Slc23a1 | XP_007904441 | Ghost shark |
| *Cyprinus carpio* | Slc23a1 | XP_018975086 | Common carp |
| *Danio rerio* | Slc23a1 | NP_001166970 | Zebrafish |
| *Gallus gallus* | Slc23a1 | XP_004944825 | Chicken |
| *Homo sapiens* | Slc23a1 | NP_005838 | Human |
| *Ictalurus punctatus* | Slc23a1 | XP_017348904 | Channel catfish |
| *Latimeria chalumnae* | Slc23a1 | XP_005989382 | Coelacanth |
| *Lepisosteus oculatus* | Slc23a1 | XP_015205203 | Spotted gar |
| *Lethenteron japonicum* | Slc23a1 | JL2986 | Japanese lamprey |
| *Monodelphis domestica* | Slc23a1 | XP_016278852 | Gray short-tailed opossum |
| *Mus musculus* | Slc23a1 | NP_035527 | House mouse |
| *Oryzias latipes* | Slc23a1 | XP_004076777 | Japanese rice fish |
| *Poecilia formosa* | Slc23a1 | XP_007556555 | Amazon molly |
| *Rhincodon typus* | Slc23a1 | XP_020385431 | Whale shark |
| *Xenopus tropicalis* | Slc23a1 | XP_002932349 | Tropical clawed frog |
| *Anolis carolinensis* | Slc23a2 | XP_003229574 | Green anole |
| *Danio rerio* | Slc23a2 | XP_021327808 | Zebrafish |
| *Gallus gallus* | Slc23a2 | XP_015152797 | Chicken |
| *Homo sapiens* | Slc23a2 | NP_976072 | Human |
| *Latimeria chalumnae* | Slc23a2 | XP_014352307 | Coelacanth |
| *Lepisosteus oculatus* | Slc23a2 | XP_015197789 | Spotted gar |
| *Lethenteron japonicum* | Slc23a2 | JL15308 | Japanese lamprey |
| *Leucoraja erinacea* | Slc23a2 | ctg10612 | Little Skate |
| *Monodelphis domestica* | Slc23a2 | XP_007476346 | Gray short-tailed opossum |
| *Mus musculus* | Slc23a2 | NP_061294 | House mouse |
| *Sarcophilus harrisii* | Slc23a2 | XP_003757995 | Tasmanian devil |
| *Scyliorhinus canicula* | Slc23a2 | ctg15352 | Small-spotted catshark |
| *Xenopus tropicalis* | Slc23a2 | NP_001120161 | Tropical clawed frog |
| *Anolis carolinensis* | Slc23a3 | XP_008108299 | Green anole |
| *Danio rerio* | Slc23a3 | XP_017213243 | Zebrafish |
| *Gallus gallus* | Slc23a3 | XP_015145565 | Chicken |
| *Homo sapiens* | Slc23a3 | NP_001138362 | Human |
| *Latimeria chalumnae* | Slc23a3 | XP_014343771 | Coelacanth |
| *Lepisosteus oculatus* | Slc23a3 | XP_015214926 | Spotted gar |
| *Monodelphis domestica* | Slc23a3 | XP_007501758 | Gray short-tailed opossum |
| *Mus musculus* | Slc23a3 | NP_919314 | House mouse |
| *Xenopus tropicalis* | Slc23a3 | XP_012826889 | Tropical clawed frog |
| *Anolis carolinensis* | Slc23a4 | XP_016849222 | Green anole |
| *Callorhynchus milii* | Slc23a4 | XP_007903282 | Elephant shark |
| *Danio rerio* | Slc23a4 | NP_001013353 | Zebrafish |
| *Gallus gallus* | Slc23a4 | XP_015145041 | Chicken |
| *Gekko japonicus* | Slc23a4 | XP_015276774 | Schlegel's Japanese gecko |
| *Latimeria chalumnae* | Slc23a4 | XP_005991386 | Coelacanth |
| *Lepisosteus oculatus* | Slc23a4 | XP_006633863 | Spotted gar |
| *Lethenteron japonicum* | Slc23a4 | JL7088 | Japanese lamprey |
| *Monodelphis domestica* | Slc23a4 | XP_007504433 | Gray short-tailed opossum |
| *Mus musculus* | Slc23a4 | XP_006506197 | House mouse |
| *Oryzias latipes* | Slc23a4 | XP_004069901 | Japanese medaka |
| *Rhincodon typus* | Slc23a4 | XP_020391140 | Whale shark |
| *Sarcophilus harrisii* | Slc23a4 | XP_012406506 | Tasmanian devil |
| *Xenopus tropicalis* | Slc23a4 | XP_012815547 | Tropical clawed frog |
| *Boleophthalmus pectinirostris* | Slc23a5 | XP_020774843 | Great blue-spotted mudskipper |
| *Cynoglossus semilaevis* | Slc23a5 | XP_016888339 | Tongue sole |
| *Cyprinus carpio* | Slc23a5 | XP_018944039 | Common carp |
| *Danio rerio* | Slc23a5 | XP_009296238 | Zebrafish |
| *Fundulus heteroclitus* | Slc23a5 | XP_021172742 | Mummichog |
| *Hippocampus comes* | Slc23a5 | XP_019750396 | Tiger tail seahorse |
| *Ictalurus punctatus* | Slc23a5 | XP_017328967 | Channel catfish |
| *Lates calcarifer* | Slc23a5 | XP_018548144 | Barramundi perch |
| *Lepisosteus oculatus* | Slc23a5 | XP_015208244 | Spotted gar |
| *Maylandia zebra* | Slc23a5 | XP_014264577 | Zebra mbuna |
| *Monopterus albus* | Slc23a5 | XP_020448997 | Swamp eel |
| *Nanorana parkeri* | Slc23a5 | XP_018412235 | - |
| *Neolamprologus brichardi* | Slc23a5 | XP_006782389 | Fairy cichlid |
| *Nothobranchius furzeri* | Slc23a5 | XP_015819289 | Turquoise killifish |
| *Oncorhynchus mykiss* | Slc23a5 | XP_021432371 | Rainbow trout |
| *Oreochromis niloticus* | Slc23a5 | XP_005454237 | Nile tilapia |
| *Oryzias latipes* | Slc23a5 | XP_004086320 | Japanese medaka |
| *Poecilia reticulata* | Slc23a5 | XP_008408845 | Guppy |
| *Pundamilia nyererei* | Slc23a5 | XP_005731628 | - |
| *Pygocentrus nattereri* | Slc23a5 | XP_017573200 | Red-bellied piranha |
| *Salmo salar* | Slc23a5 | XP_013981136 | Atlantic salmon |
| *Scleropages formosus* | Slc23a5 | XP_018602707 | Asian bonytongue |
| *Sinocyclocheilus rhinoceros* | Slc23a5 | XP_016366936 | - |
| *Stegastes partitus* | Slc23a5 | XP_008289900 | Bicolor damselfish |
| *Takifugu rubripes* | Slc23a5 | XP_003967776 | Fugu rubripes |
| *Xenopus tropicalis* | Slc23a5 | XP_002937113 | Tropical clawed frog |

**Table S2.**
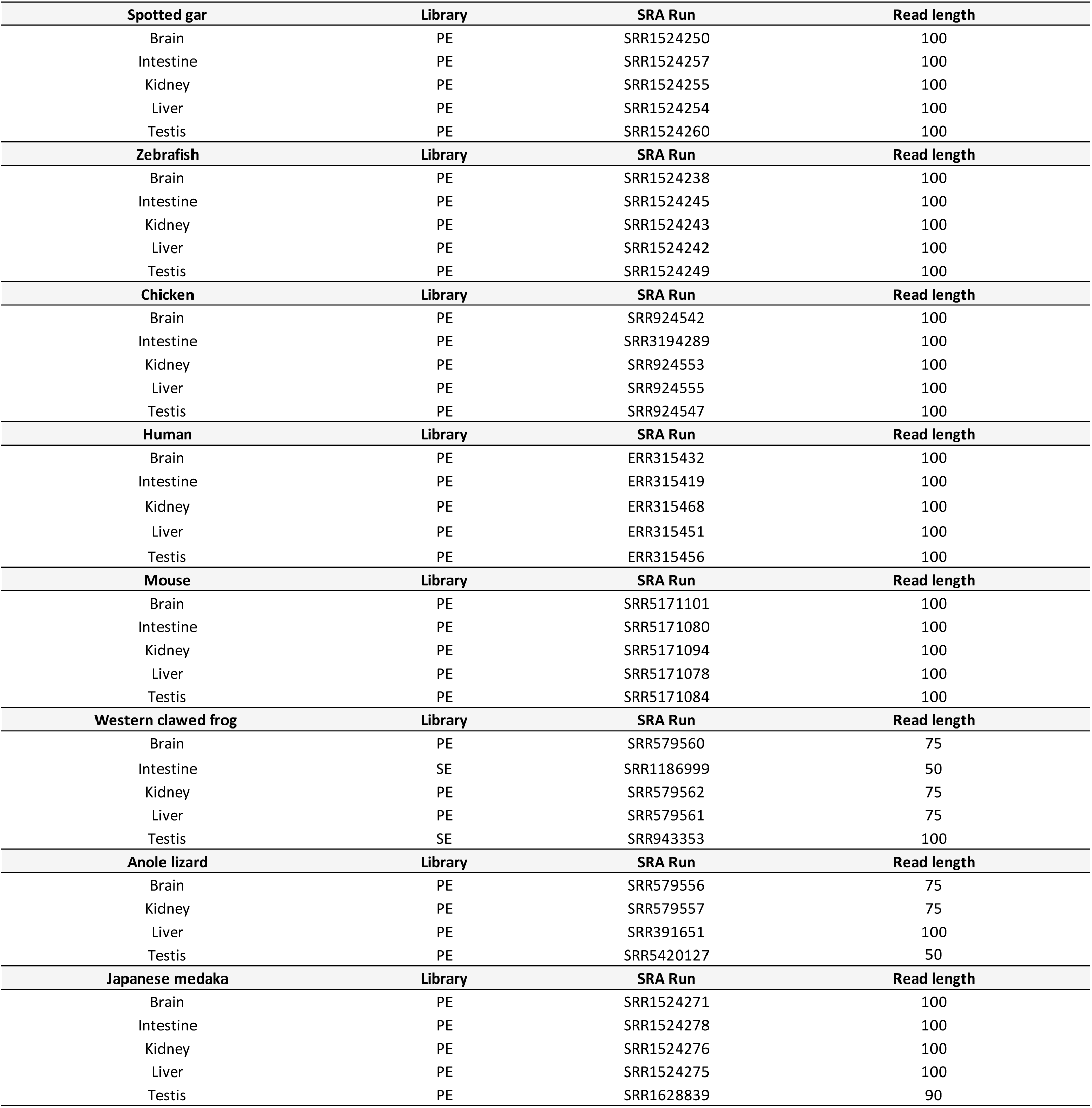
Accession numbers of the RNAseq files. PE – portable executable format.

| Spotted gar | Library | SRA Run | Read length |
| --- | --- | --- | --- |
| Brain | PE | SRR1524250 | 100 |
| Intestine | PE | SRR1524257 | 100 |
| Kidney | PE | SRR1524255 | 100 |
| Liver | PE | SRR1524254 | 100 |
| Testis | PE | SRR1524260 | 100 |
| Zebrafish | Library | SRA Run | Read length |
| Brain | PE | SRR1524238 | 100 |
| Intestine | PE | SRR1524245 | 100 |
| Kidney | PE | SRR1524243 | 100 |
| Liver | PE | SRR1524242 | 100 |
| Testis | PE | SRR1524249 | 100 |
| Chicken | Library | SRA Run | Read length |
| Brain | PE | SRR924542 | 100 |
| Intestine | PE | SRR3194289 | 100 |
| Kidney | PE | SRR924553 | 100 |
| Liver | PE | SRR924555 | 100 |
| Testis | PE | SRR924547 | 100 |
| Human | Library | SRA Run | Read length |
| Brain | PE | ERR315432 | 100 |
| Intestine | PE | ERR315419 | 100 |
| Kidney | PE | ERR315468 | 100 |
| Liver | PE | ERR315451 | 100 |
| Testis | PE | ERR315456 | 100 |
| Mouse | Library | SRA Run | Read length |
| Brain | PE | SRR5171101 | 100 |
| Intestine | PE | SRR5171080 | 100 |
| Kidney | PE | SRR5171094 | 100 |
| Liver | PE | SRR5171078 | 100 |
| Testis | PE | SRR5171084 | 100 |
| Western clawed frog | Library | SRA Run | Read length |
| Brain | PE | SRR579560 | 75 |
| Intestine | SE | SRR1186999 | 50 |
| Kidney | PE | SRR579562 | 75 |
| Liver | PE | SRR579561 | 75 |
| Testis | SE | SRR943353 | 100 |
| Anole lizard | Library | SRA Run | Read length |
| Brain | PE | SRR579556 | 75 |
| Kidney | PE | SRR579557 | 75 |
| Liver | PE | SRR391651 | 100 |
| Testis | PE | SRR5420127 | 50 |
| Japanese medaka | Library | SRA Run | Read length |
| Brain | PE | SRR1524271 | 100 |
| Intestine | PE | SRR1524278 | 100 |
| Kidney | PE | SRR1524276 | 100 |
| Liver | PE | SRR1524275 | 100 |
| Testis | PE | SRR1628839 | 90 |

**Table S3.**
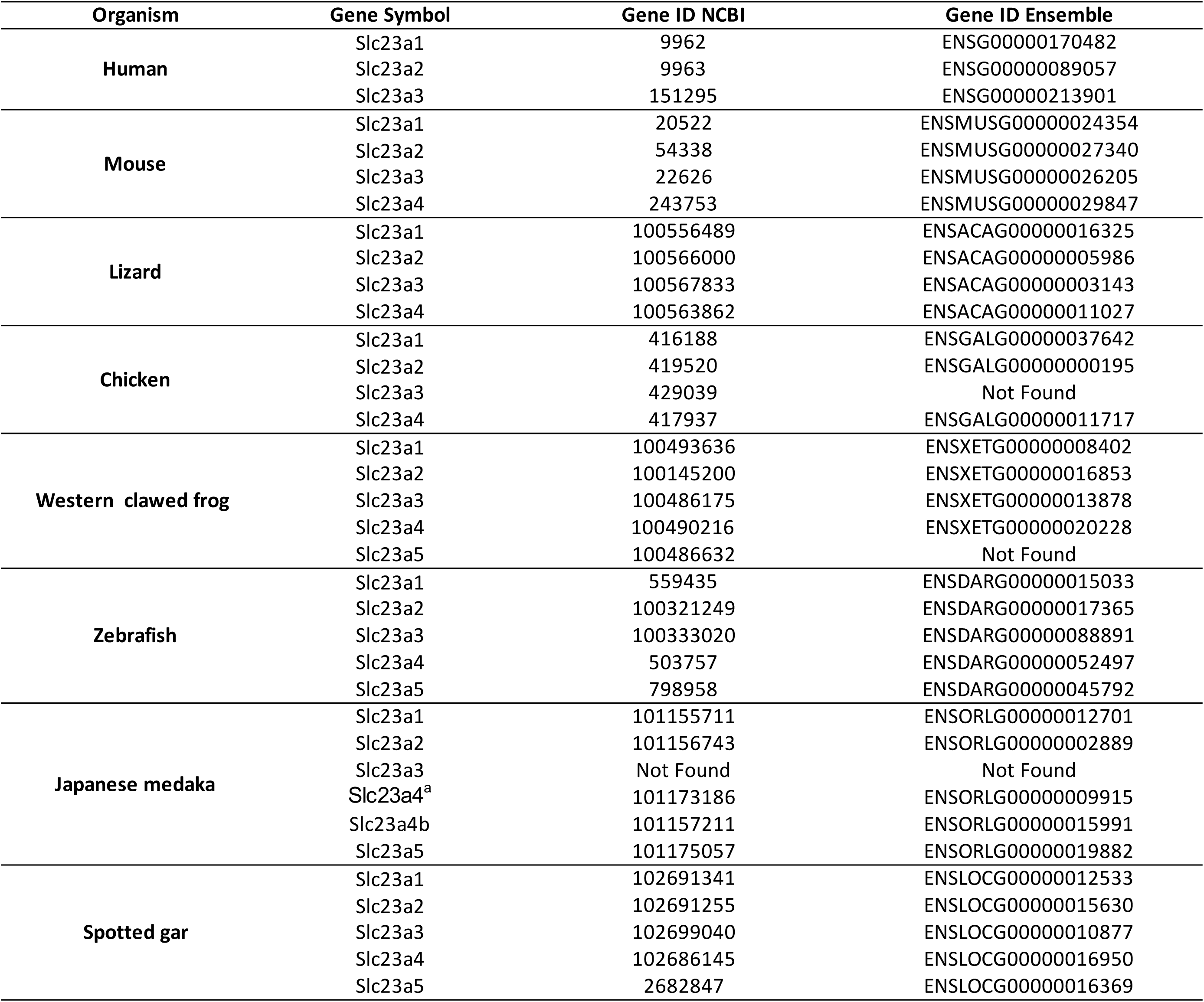
Gene identifier for all genes included in gene expression analysis, collected from NCBI and ENSEMBLE Databases.

**Table S4.**
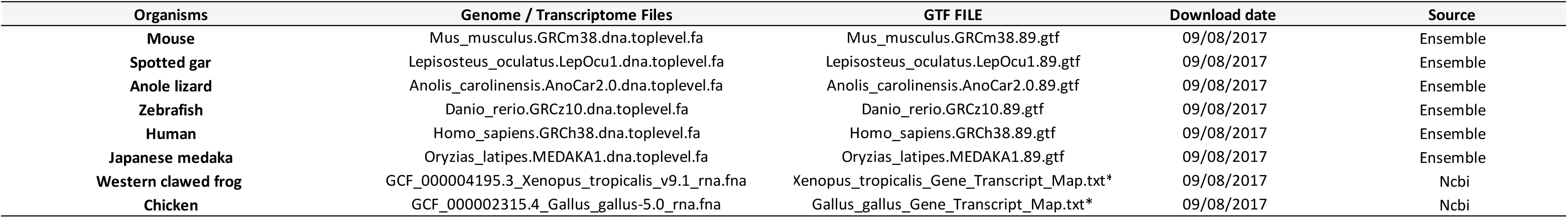
Genome and GTF files retrieved from Ensemble database (Release 89) and Transcriptome files retrieved from NCBI used for this study.

**Table S5.** Relative gene expression levels of Slc23 family genes for eight species. The values are presented in log2 (TPM +1) ratios.

| Organism | Tissue | Slc23a1 | Slc23a2 | Slc23a3 | Slc23a4a | Slc23a4b | Slc23a5 |
| --- | --- | --- | --- | --- | --- | --- | --- |
| Human | BRAIN | 0,00 | 5,82 | 0,00 | - | - | - |
|  | INTESTINE | 5,85 | 2,64 | 4,61 | - | - | - |
|  | KIDNEY | 3,71 | 3,70 | 4,28 | - | - | - |
|  | LIVER | 4,03 | 3,58 | 2,36 | - | - | - |
|  | TESTIS | 0,00 | 4,59 | 0,00 | - | - | - |
| Mouse | BRAIN | 0,00 | 4,91 | 0,00 | 0,00 | - | - |
|  | INTESTINE | 3,73 | 5,24 | 1,02 | 5,89 | - | - |
|  | KIDNEY | 6,91 | 2,82 | 4,08 | 0,00 | - | - |
|  | LIVER | 5,06 | 2,85 | 0,00 | 0,00 | - | - |
|  | TESTIS | 0,00 | 1,63 | 0,00 | 0,00 | - | - |
| Lizard | BRAIN | 0,91 | 2,82 | 0,72 | 2,49 | - | - |
|  | KIDNEY | 6,76 | 1,39 | 6,10 | 1,19 | - | - |
|  | LIVER | 3,02 | 0,00 | 0,00 | 0,65 | - | - |
|  | TESTIS | 0,00 | 0,00 | 0,00 | 0,00 | - | - |
| Chicken | BRAIN | 0,00 | 6,27 | 0,96 | 1,82 | - | - |
|  | INTESTINE | 6,44 | 3,25 | 2,96 | 6,09 | - | - |
|  | KIDNEY | 7,78 | 1,49 | 4,17 | 1,60 | - | - |
|  | LIVER | 5,20 | 1,98 | 1,76 | 0,82 | - | - |
|  | TESTIS | 2,35 | 2,08 | 2,03 | 1,85 | - | - |
| Western clawed frog | BRAIN | 0,00 | 6,45 | 0,00 | 2,54 | - | 0,00 |
|  | INTESTINE | 0,00 | 2,23 | 0,00 | 5,65 | - | 0,00 |
|  | KIDNEY | 7,63 | 4,08 | 6,53 | 5,00 | - | 5,40 |
|  | LIVER | 0,00 | 3,41 | 0,00 | 2,14 | - | 0,00 |
|  | TESTIS | 2,69 | 4,01 | 0,00 | 1,49 | - | 1,64 |
| Zebrafish | BRAIN | 0,85 | 3,73 | 0,71 | 0,00 | - | 1,01 |
|  | INTESTINE | 2,69 | 1,21 | 0,00 | 4,00 | - | 0,00 |
|  | KIDNEY | 5,35 | 2,27 | 3,42 | 4,78 | - | 0,93 |
|  | LIVER | 0,00 | 1,26 | 0,00 | 0,00 | - | 0,00 |
|  | TESTIS | 0,93 | 2,74 | 0,00 | 0,63 | - | 1,97 |
| Japanese medaka | BRAIN | 0,00 | 4,70 | - | 0,00 | 0,00 | 0,84 |
|  | INTESTINE | 2,76 | 2,91 | - | 4,95 | 0,00 | 0,00 |
|  | KIDNEY | 3,89 | 3,02 | - | 4,43 | 0,00 | 0,00 |
|  | LIVER | 0,00 | 4,06 | - | 0,00 | 0,00 | 0,00 |
|  | TESTIS | 0,00 | 3,20 | - | 0,60 | 0,00 | 0,00 |
| Spotted gar | BRAIN | 0,58 | 4,94 | 0,00 | 0,00 | - | 0,00 |
|  | INTESTINE | 5,67 | 0,96 | 0,00 | 0,00 | - | 0,00 |
|  | KIDNEY | 3,18 | 0,90 | 1,78 | 0,86 | - | 0,00 |
|  | LIVER | 0,74 | 0,78 | 0,00 | 0,00 | - | 0,00 |
|  | TESTIS | 1,92 | 7,21 | 0,00 | 0,00 | - | 0,00 |

**Table S6.** Renal uric acid absorption and secretion: the urate transportome. n.f. – not found.

|  | Transporter | *H. sapiens* | *M. musculus* | *G. gallus* | *A. carolinensis* | *X. tropicalis* | *D. rerio* | *L. oculatus* |
| --- | --- | --- | --- | --- | --- | --- | --- | --- |
| **Basolateral Absorption** | SLC2A9/GLUT9 | ✓ | ✓ | ✓ | ✓ | ✓ | ✓ | ✓ |
| **Apical Absorption** | SLC22A11/OAT4 | ✓ | n.f. | n.f. | n.f. | n.f. | n.f. | n.f. |
|  | SLC22A12/URAT1 | ✓ | ✓ | n.f. | n.f. | n.f. | n.f. | n.f. |
|  | SLC22A13/OAT10 | ✓ | ✓ | ✓ | ✓ | ✓ | ✓ | ✓ |
| **Basolateral Secretion** | SLC22A6/OAT1 | ✓ | ✓ | ✓ | ✓ | ✓ | n.f. | n.f. |
|  | SLC22A8/OAT3 | ✓ | ✓ |  |  |  | n.f. | n.f. |
| **Apical Secretion** | SLC17A1/NPT1 | ✓ | ✓ | n.f. | n.f. | n.f. | n.f. | n.f. |
|  | SLC17A3/NPT4 | ✓ | ✓ | n.f. | n.f. | n.f. | n.f. | n.f. |

